# Impact of abiotic conditions on Fungal Diversity and Comparative analysis of Soil quality of two distinct locations

**DOI:** 10.1101/2022.09.30.510421

**Authors:** Sindhushri Chauhan, Parashuram Ganure, Chethan J. Dandin

## Abstract

Soil fertility play a prominent role in biological sustenance in turn fungi are significant participants in bridging biotic and abiotic interactions between the components in the environment, and they involve in breakdown of organic/ inorganic materials during bioleaching/ bioconversion leading to bio-availability and thus recycling of nutrients. Local environmental factors (abiotic), such as the chemical and physical properties of the soil, greatly influence the composition of existing fungal communities and determine the diversity of soil fungi. A correlation study of the micro fungi found in ecologically two different regions (Karnataka state, India) of Uttara Kannada and Chitradurga districts. One being part of India’s Western Ghats, a biologically fertile ecosystem and the other being a dry, arid region of Deccan plateau with scanty rain fall. Soil samples were collected from various geographical locations in order to study and evaluate the role that certain soil physicochemical features play in fungi diversity. According to our findings, soil attributes (fertility) are related to the composition, richness, and diversity of fungi in soils and thus soil’s physico-chemical properties assessed were linked to fungal diversity measurements. This led us to indicate the role of abiotic factors effecting the fungal diversity in soil and their prominent role in soil fertility.

**Importance of the Work:** This work signifies the role of fungal diversity on soil fertility and the impact of abiotic and physico-chemical soil parameters on the same. Thus the work explores the importance of the each of these components and the wild fungal strain’s participation in value addition to their functional role in breakdown of organic/ inorganic materials during bioleaching/ bioconversion leading to bio-availability and thus recycling of nutrients. The work also highlights soil pH as one of key indicators to define the diversity of soil fungi and can provide a direct correlation between pH, bioconversion and fungal activity/ diversity. This work is first of a kind from southern India connecting the western ghat bio-diversity hot-spot fertile region with the neighbouring barren Bayaluseemae region of Karnataka State, South India with regards to abiotic factors, fungi and fertility.

## 1. Introduction

The presence and bioavailability of various nutrients such as nitrogen, phosphorus, and potassium for microbial activity in the soil contribute to the richness of soil fertility. Soil fertility is determined by three interconnected and mutually dependent factors: physical fertility, chemical fertility, and biological fertility. Among the three Biological fertility refers to the microorganism’s role and decomposition of organic matter that interact with other soil components. According to a recent study on the diversity of soil fungi, there are approximately 80,500 operational taxonomic units (OTUs) of fungi found in soils worldwide (Tedersoo, *et al*., 2014). Some of them are well-known for their contributions to nutrient cycling and biogeochemical cycles. The biological-microbial component is the least thoroughly evaluated and understood of the three fertility components, particularly fungi and the rich diversity of others such as viruses, bacteria, protozoans, and algae that are part of the most complex, interactive microbial communities.

Microbial biotechnology through the exploration of soil microbial resources and diversity has proven to be one of the most powerful and potent tools that could provide insights into nutrient limitations (notably N, P and K) and their recycling in most of the soils.

Recent research has demonstrated that soil pH is the primary factor influencing microbial richness, but other soil characteristics, such as micro and macro components, can also affect the community’s composition and variety (Sun and Zhalnina, *et al*., 2015). It has also been discovered that soil microbial populations are significantly influenced by the availability of soil nutrients both organic and inorganic nutrients, thus also been demonstrated that the organisation of soil microbial communities significantly correlates with soil moisture and salinity. In some particular habitats, studies have indicated that climate change effects increase in CO_2_, temperature, and precipitation that can affect the overall makeup of soil microbial communities (Rinnan, 2009 and Chen, *et al*., 2015) such variety of factors can influence soil microbial communities, but the driving factor may at times differ between ecosystems. Most soil microbial diversity studies have been conducted in natural settings, so baring wild properties, but urban ecosystems are more complex than other ecosystems due to unusual anthropogenic activities (Civerolo, *et al*., 2007 and Ash, *et al*., 2008). It is hypothesised that changes in soil chemical and physical properties may have an impact on soil microbial diversity in urban areas. The impact of urbanisation on soil microbial communities has gotten little attention. Notably, the current study shows that physico-chemical characteristics of the soil are mostly connected with fungal diversity and community composition. Numerous earlier research have demonstrated the importance of soil pH in determining the makeup and characteristics of the soil microbial populations. One of the simplest drivers of variance in microbial diversity is thought to be soil pH, which is consistently accurate predictor of the patterns of fungal distribution, but is not the only one. Fungi may survive in a wide range of pH (3.0 to 10.0), temperature, and habitats, including both fertile and non-fertile environments (0 to 90 °C) (Franc *et al*., 2015). Microbiologists and agriculturalists have long been very interested in understanding the chemical and physical properties of soil to assess the level of soil fertility, which attributes as effectors involving microbial diversity for this change. Our aim in the present study is to investigate, analyse and correlate the diversity of terrestrial soil fungi along two different climatic zones typically correlating their participatory roles in respective environmental conditions for bioavailability and soil fertility as an important function. The screening and isolation, was from selected two regions of which one belonging to Uttara Kannada (forest with high soil fertility and bioconversion rate) and other being barren or eroded (lands that doesn’t support agriculture due to lack of natural soil fertility and rain) in Chitradurga district. Microbial diversity in barren soil samples as well as fertile soil samples are largely attributed to natural environment and its ability to adapt to the environmental stress factors such as low moisture availability and low or high temperature, minimum available moisture in most of the barren lands that is influenced by limited precipitation in the form of rainfall and fog derived moisture (Warren-Rodes *et al*., 2006). The emphasis of the study is about microbial diversity and its wild behaviour to various stress factors ranging from very low moisture and high temperature of Chitradurga district and on contrary soil with high fertility and bioconversion rate of Uttara Kannada district.

## 2. Materials and procedures

### 2.1. Location study and Sampling

Like air and water, soil is an important natural resource. There are many different types of soil in the Indian subcontinent, and the altitude, climate, disproportionate rainfall, mineral composition, biodiversity, and many other related factors all have an impact on how those soils are formed. Different regions of the country have different types of soil. A century ago, the soil was categorised according to its fertility, but today, a variety of factors and their bioavailability are taken into account, and so soil types are categorised according to their pH, colour, OC percentage, moisture, N, P, K content, soil biodiversity, etc.

To characterize and compare the fertility status of soil, samples were collected from two different ecosystems, both the regions are geographically distinct one is in Uttara Kannada district which consists of highly fertile forest region (majorly undisturbed soil ecosystems), another one is from the Chitradurga district with highly fertility-eroded which is commonly called as Bayaluseme (in Kannada), disturbed region i.e., lands not under use for agriculture due to lack/ loss of fertility (Fig. 1).

**Fig.1:**
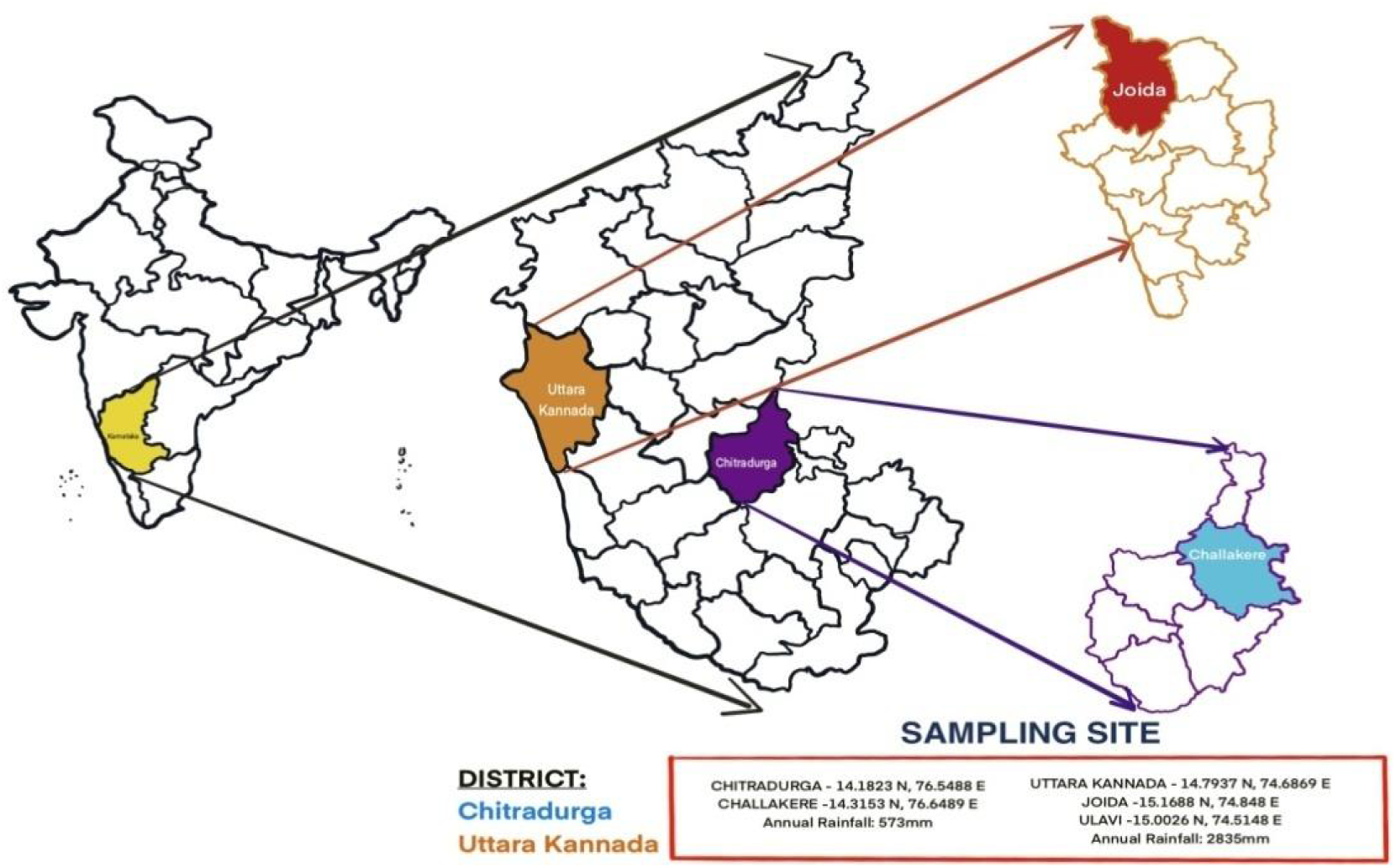
Map depicting the two locations Challakere Tq of Chitradurga Dist. and Joida Tq of Uttara Kannada Dist. of Karnataka state from where the comparative soil samplings were done for the present study.

Karnataka is climatically pounded with some of the most magnificent tropical forests of the Indian subcontinent. Different region of land mass have different soil types and are categorised according to its soil fertility factors (pH, color, OC%, etc.) The Western Ghats, is among the world’s major biodiversity hotspots, are abundant in this state and come in many different varieties with enormous forest biodiversity of, about 60% of Karnataka’s forest are situated. Western Ghats of **Uttara Kannada** district is known for their dense and undisturbed natural forest area which covers about 80% of the total area of the district.

On the contrary, **Chitradurga**, which covers an area of 8,440 square kilometres, is one of the drought-prone districts of Karnataka state in southern India. The soil texture is partly black and mixed red/black soil, red loamy and sandy soil, with low to moderate rainfall and no rain-fed regions. The entire district is located in the valley of the Krishna and Vedavathi River basins, with the Tungabhadra River flowing northwest from Chitradurga.

### 2.2. Collection of soil samples and their physico-chemical analysis

Site-specific and random sampling methods are examples of soil sampling strategies. In the current study, samples were collected at a depth of 10-15cm from each zone using a random sampling methodology after scraping the top 1-2cm layer of soil. The soil samples were collected in sealed polythene bags and transported in a closed container to the laboratory. Standard protocols were used to determine the physicochemical properties of soil samples (Kekane, 2015). The physico-chemical studies of soil parameters being an important guideline to agricultural chemists ascertain them in assessing the soil fertility, bioavailability and thereby plant growth, adaptation and also in soil bio-fertility management. Soil fertility has been attributed to the microbial distribution, diversity and in particular soil fungi. The soil food web is dominated by fungal populations (although they are less in number than the bacteria). Fungi can effectively assimilate 40–55% of carbon soil organic matter (SOM) source, thus are more efficient and so they can store and recycle more carbon (C) compared to bacteria.

### 2.1. Isolation of fungi

Bio-fertility is an important component of soil fertility management, among which are the microbial communities such as soil fungi, bacteria and other microorganisms that can provide exceptional value addition to the bio-conversion and bio-availability of nutrients (Briones, 2018, Snow, 2020 and Mayer *et al*., 2021). Based on the following method, outlined by Rosas-Medina and Piepenbring (2018), the soil fungi were also isolated in the current study using a small amount (approximately 0.05g) of various soil samples on sterile media such as PDA, CZA, YPDA, and SDA, respectively. The technique is a variation on Warcup soil plate technique, in which a suitable amount of soil in water suspension is spread on the surface of the agar medium. The plates were incubated in an incubation chamber at 25°C for up to 7 days or until colonies formed. This procedure entailed daily observation of culture plates, making dilutions in water to separate spores, and re-culturing until pure cultures were obtained. Later, the purely grown fungal colonies were identified morphologically and microscopically to the nearest possible taxonomic identity referring to the fungal identification key manual (Watanabe, 2002).

### 2.2. Fungal diversity index

From two location specific soil samples, fertile and barren, the number of soil samples taken, the number of fungal colonies isolated, and the number of colonies of individual fungus were recorded. Simpson’s Diversity index (D) was applied to calculate their diversity indices: which is defined as simple mathematical derivation that measures number of species present and relative abundance of each species’s diversity in a community and is measured as D, that ranges between 0 and 1, ‘0’ being infinite diversity and ‘1’ being biodiversity. It is calculated using the formula as below. Simpson’s scale D= 1-[ni (ni-1)/N (N-1)].

Where ‘ni’ is the abundance of individual species.

N = Total number of organisms.

The D has a value between 0 and 1.

D = 0 represents the infinite diversity, while D = 1 represents no diversity.

### 2.3. Identification of soil fungi

Fungal colonies were grown and isolated on agar media at 25 to30 °C for three to seven days or until colonies were observed and further used to study colony and microscopic morphology. True fungal mycelia bearing spores and unicellular cells were stained with lacto-phenol cotton blue, cautiously distributing the biomass on a microscopic slide, covering with coverslips, was observed under 10X, 40X and 100X magnifications of compound microscope. Identification of the isolated fungal colonies up to genus level was accomplished through visual inspection of morphology and structures using bright field microscopy (Aneja 2006) and using reference key manual - Pictorial Atlas of Soil and Seed Fungi (Watanabe, 2010).

## 3. Results

### 3.1. Physico-chemical characteristics of the soil

A balanced soil pH and EC correlated with maximal nutritional availability, EC was normal in both the regions i.e <0.8 ds/m. There was a significant difference found in the pH of soil collected from two geographically distinct locations, the pH of soil collected from dense/ undisturbed forest region was found to be acidic (5.067) which is conducive for rich diversity of fungi in the soil, whereas pH recorded in barren soil was found to be neutral to alkaline with the mean value of 8.11 indicating minimal diversity, supporting soil fungi.

The % OC in forest region is higher than the barren/undisturbed region, this is due to the presence of organic waste decomposed in the forest which gets recycled and adds more organic matter to the soil. The N, P, K contents were present in adequate amount in both the regions and their respective composition is shown in table 1. The graph in fig. 2 represents the comparison between physico-chemical properties of Uttara Kannada and Chitradurga soil samples. The above results (table 1) shows that the physico-chemical parameters (pH, EC, % OC, N, P, K) are directly proportional to the bioavailable nutrients resulting in rich fungal diversity in soil of Western ghats, collected from Uttara Kannada region.

**Fig.2:**
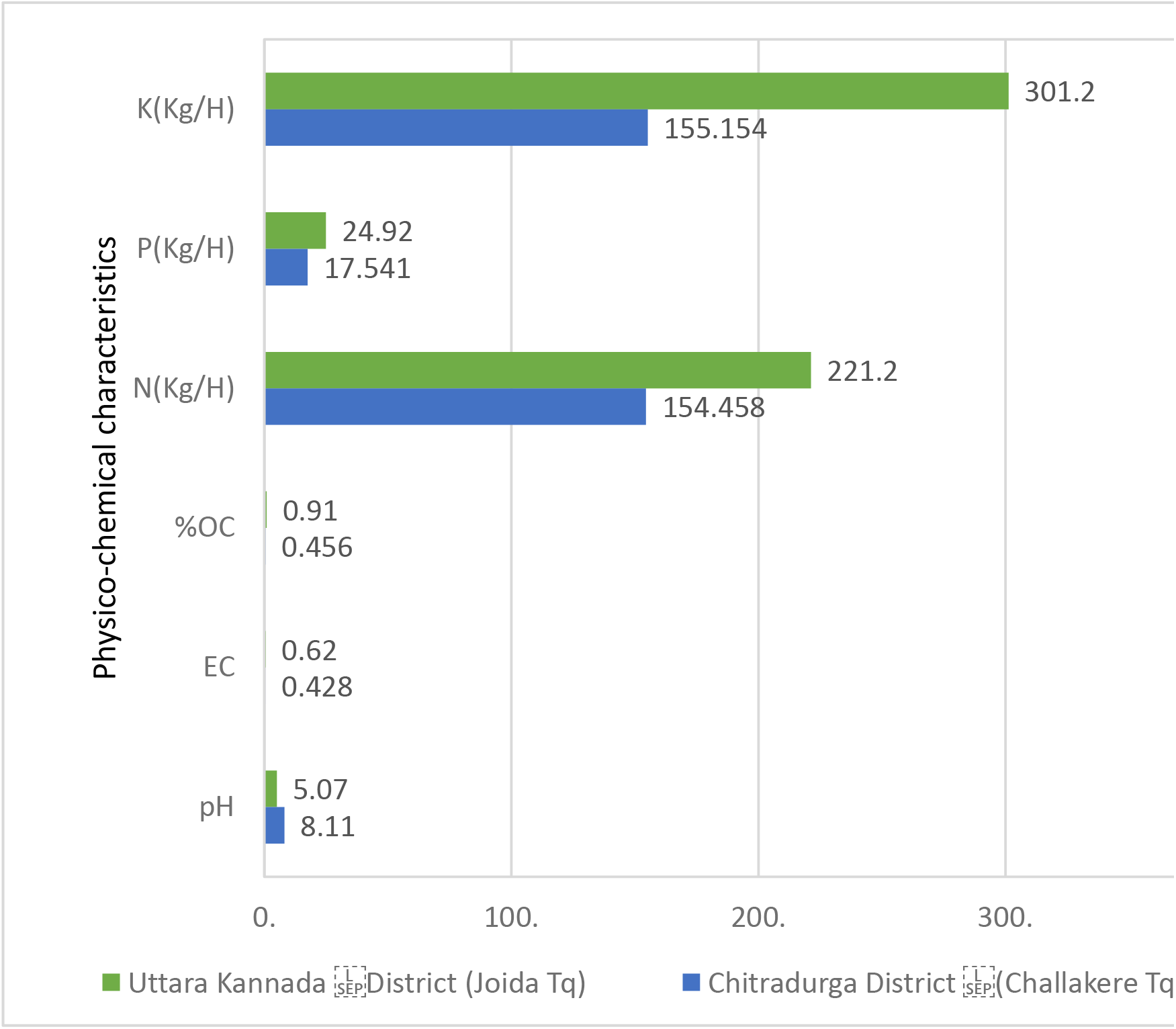
Graphical representation of comparative physico-chemical parameters of soil samples collected from Uttara Kannada and Chitradurga districts sampling site.

**Table 1:** Statistical comparison of the soil physico-chemical parameters collected across two locations by one way ANOVA. The standard deviation (n =10) mentioned with the average values of parameters. Tukey’s test was used to differentiate the mean values at a significance level of p<0.05 (mean ±SD). ^a^ & ^b^ demotion are Tukey’s test showing that there is significant difference at p <0.05. Statistical analysis was carried out using Origin-9 software (Origin Lab Corporation, Massachusetts, USA) and by one way analysis of variance (ANOVA).

| Regions | pH | EC | %OC | N(Kg/H) | P(Kg/H) | K(Kg/H) |
| --- | --- | --- | --- | --- | --- | --- |
| Uttara Kannada district (Joida Tq) | 5.067<br>±0.102<sup>b</sup> | 0.622<br>±0.193<sup>a</sup> | 0.908<br>±0.255<sup>a</sup> | 221.2<br>±5.212<sup>a</sup> | 24.92<br>±2.295<sup>a</sup> | 301.2<br>±31.923<sup>a</sup> |
| Chitradurga district (Challakere Tq) | 8.11<br>±0.54<sup>a</sup> | 0.428<br>±0.285<sup>a</sup> | 0.456<br>±0.179<sup>b</sup> | 154.458<br>±77.3<sup>b</sup> | 17.541<br>±13.117<sup>b</sup> | 155.154<br>±117.727<sup>b</sup> |

### 3.2. Isolation of fungi

In the present investigation the diversity of soil fungi was identified by using suitable media by crowd plate method along different climate zones. From both the districts 20 samples were screened for the fungal diversity and about 80 fungi were identified by using relevant key literature. Among them *Penicillium* and *Aspergillus* spp. were prominent followed by *Trichoderma, Fusarium, Cladosporium, Rhodotorula* etc. Previously, numerous studies have sought to understand how the vast, diverse soil microorganisms are influenced by abiotic and biotic factors. In the current study, forest zones had higher levels of fungal diversity than barren soil, which may be related to variations in abiotic variables. As shown in Fig 4 every single colony of different shapes and colours appeared on the plate after the first incubation was sub-cultured and maintained on another new agar plate.

**Fig 3:**
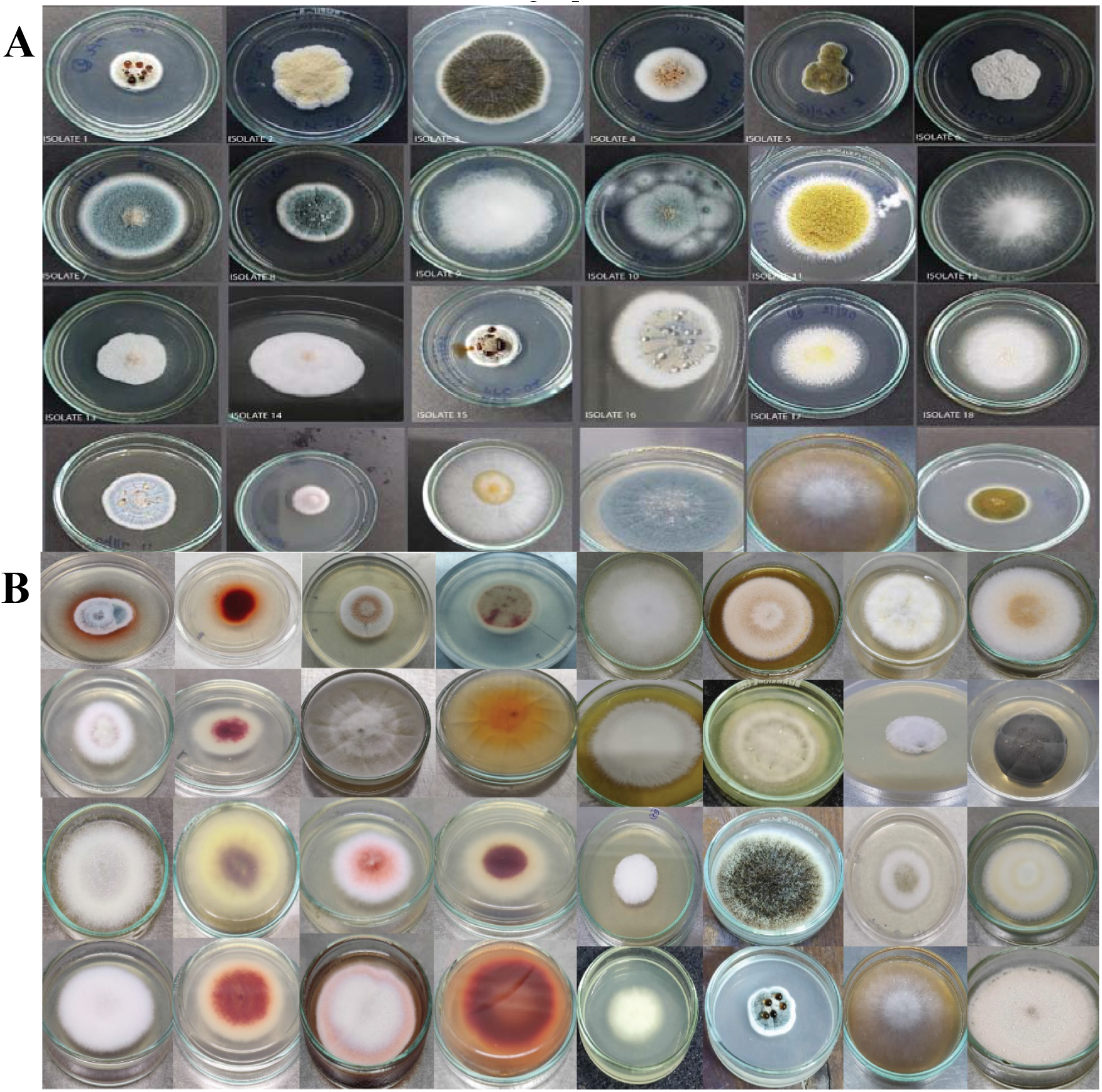
Pure cultures of different fungal isolates from Uttara Kannada (D) of Joida (T) forest region (Plate A & B).

**Fig 4:**
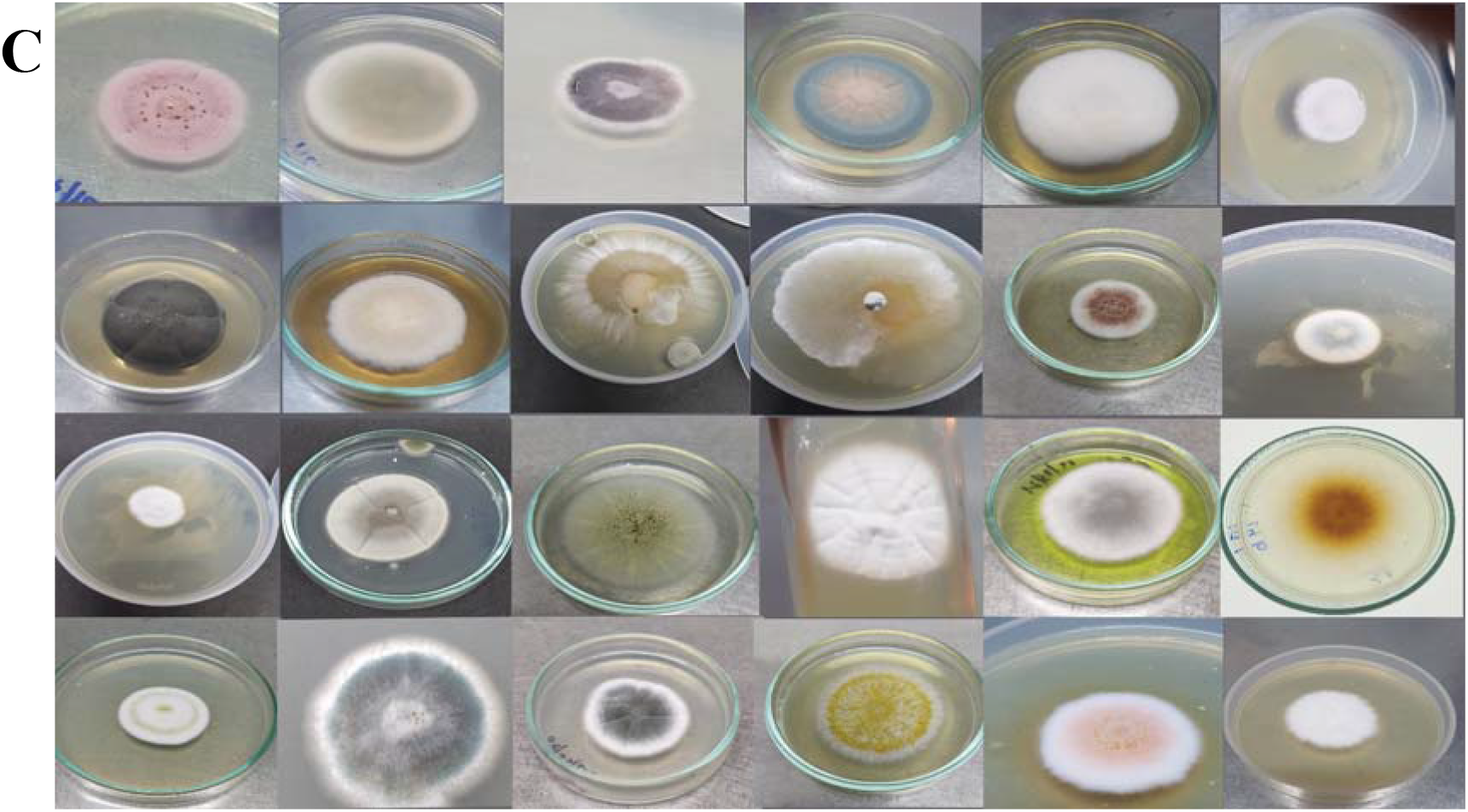
Pure cultures of different fungal isolates from Chitradurga (D) of Challakere (T) barren region (Plate C).

### 3.3. Fungal diversity index

Diversity in contrast to arid (0.1-diversity index) soil, fig 4 forest soil has a Simpson index of 0.90. Low scores (near 0) indicate low diversity (Table 2), whereas high scores (near 1) suggest high diversity, which implies that the forest soil contains the greatest diversity of fungi. Fungi are benefited most, from a variety of nutrients in forest soil and therefore, the greatest variety of fungus can grow in this soil, suggesting that the fungal diversity in arid soil is low.

**Table 2:**
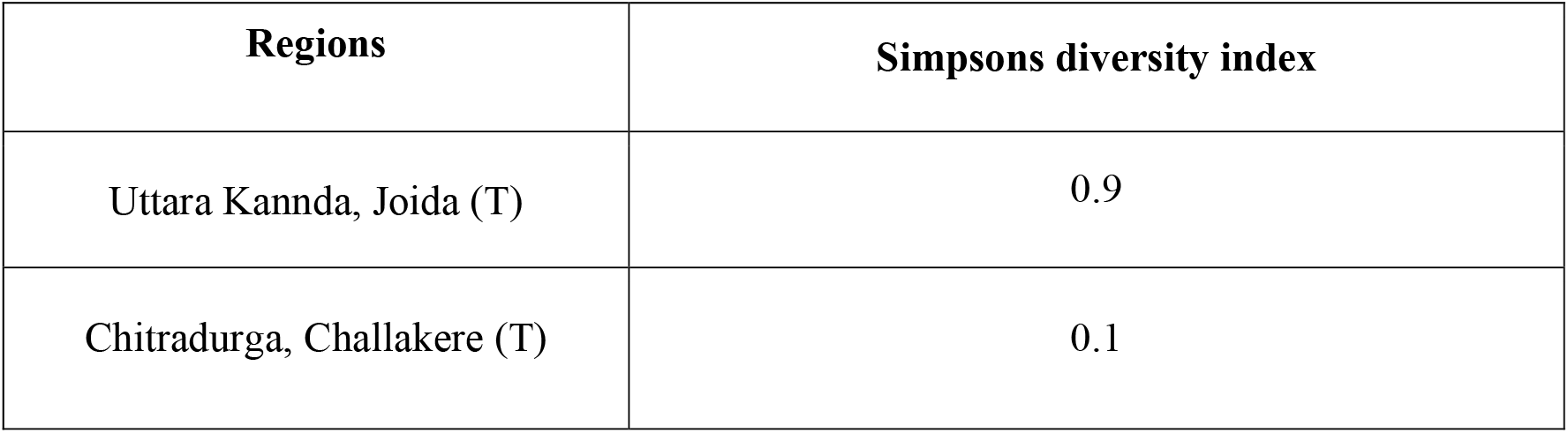
Simpson’s diversity index

**Fig 4:**
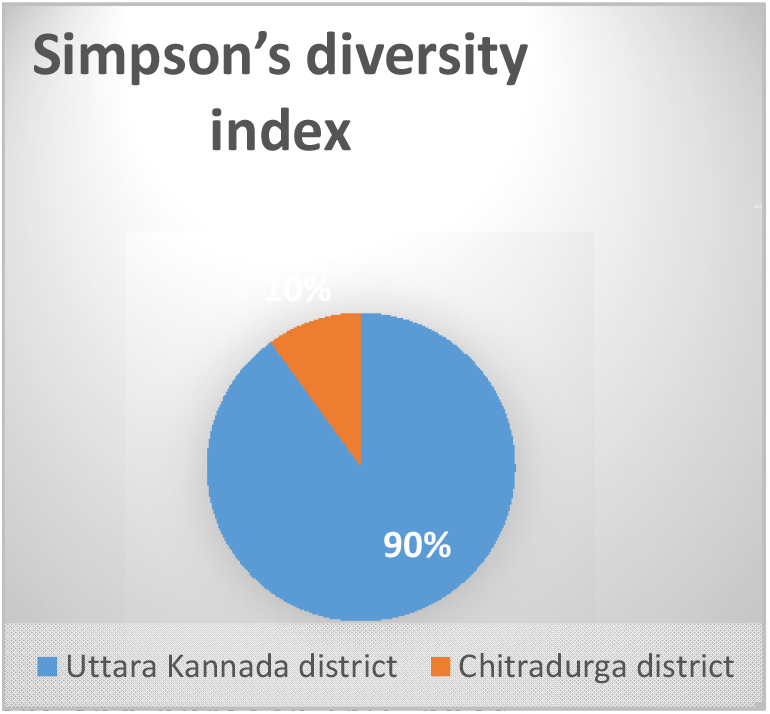
Comparative pie chart of relative percentage of Simpson’s diversity index.

The ability of microbial colonisation in barren soil, against environmental stress impact, such as limited moisture availability, low or high temperature and UV radiation has evaluated in detail. Fungal cultures obtained from various isolation techniques were used with a number of permutation and combination that benefits the chances of maximum isolation. In the current study to investigate the microbial reservoirs of barren soil samples. During our investigation, we could isolate a wide range of fungal and yeast strains, many of which were previously assumed to be present in smaller numbers.

## 4. Discussion

In this work, the soil sampling has provided an inclusive information about the biological species diversity particularly of the fungal isolates playing effective role in soil fertility. The first indications are with the pH of the soil that consistently was around 5.0 (mean average ±5.607) which is indicative of efficient fungal colonization as it’s the nominal growth pH of predominantly fungal species than the bacterial growth and activity (Rousk *et al*., 2009). There was a clear difference in the % OC (organic carbon) content and nitrogen (N), phosphorus (P) and potassium (K) by more than 40% in total giving an additional indication of the function of fungal diversity effectively playing the role in overall enhancement of the soil fertility index. Other researchers have pointed to the physico-chemical features of soil, including Dasgupta and Brahmaprakash, (2021), who studied and analysed the soil microorganisms’ response to abiotic variables, this further contributing to variation in soil nutrient availability. Additional factors that are not directly related to the specific environmental conditions were not assessed in this work, and these additional factors may contribute to the diversification of microbial communities.These variables could include: the purposeful movement of microorganisms independently and associated species, through the soil profile by water or bioturbation (Lavelle et al., 1997, Young and Crawford, 2004) causing change in geochemistry (Richter and Markewitz, 1995), organic composition of the soil (Lal, 2004, Johnson et al., 2005; Balkwill *et al*., 1998 and Kieft *et al*., 1998).

The concentration of hydrogen ions (H^+^) in the soil solution a factor attributed to the microbial activity of biodegradation and mineral leaching, determines the pH of the soil, there by indicates how acidic or alkaline is the soil. Lower pH values are associated with higher H^+^ ion concentrations, and vice versa (Rousk *et al*., 2009). The importance of pH in preserving soil fertility cannot be overstated (Patil *et al*., 2014). Electrical Conductivity/Conductance (EC) is a positive indicator for crops since it aids in nutrient absorption (Martin *et al*., 2011). Low EC values are shown to be suitable for plant growth, indicating better fertility (Jain *et al*., 2015). One of the important chemical indicators of soil quality is the amount of soil organic matter, often known as soil organic carbon. It influences soil porosity and fosters water and gas exchange processes. Our findings demonstrated that the levels of OC in the soil samples from Chitradurga and Uttara Kannada differed significantly and were much greater when compared. Soil OM has a significant function to play in preserving the physical, chemical, and biological qualities of the soil as well as the crop productivity and output.

The status of the available organic carbon may be connected to the variation in N content. Low organic matter content, little rainfall, and little vegetation in these soils all contribute to faster decomposition and removal of organic matter, which results in low nitrogen status. According to (Anon, 1998), 35.8% of the soils in the state of Karnataka had moderate nitrogen availability, especially in the plateau’s hilly regions and irrigated areas, while the remaining soils had low nitrogen availability overall.

When comparing soil samples from forests and barren areas, it was discovered that the amount of potassium in the former was substantially higher. There was an adequate supply of potassium in both areas. An increase in K might be the result of soil saturation, which widened clay minerals and released previously fixed K, as well as huge fertiliser storage, which caused these to dissolve in floodwater. A higher osmotic pressure in the plant is caused by more potassium in the soil, which increases its ability to absorb water (Joseph, 2005).

Phosphorus is a limiting nutrient that is found in plant nuclei and functions as a kind of energy storage (Jain *et al*., 2014). Phosphorus levels in soil samples from forests in both zones were noticeably higher than those from arid areas. Greater organic matter concentration may be the cause of higher phosphorus levels. Compared to soils with low organic content, soils rich in organic matter provide plants with organic phosphates (Miller and Donahne, 2001). In contrast to soils with maximal leaching, soils with less leaching impact have higher levels of phosphorus (Ashraf *et al*., 2012). There are certain methodological restrictions on this study. Due to the fact that many species are difficult to identify on agar media, the isolation of fungi in culture is not complete, some species grow faster than others when soil particles are distributed over the medium’s surface due to their rapid growth rates (Kirk *et al*., 2008). In which case the higher fungal species diversity value would be 0.90 of Joida forest of Uttara Kannada with barren soil from challakere region of Chitradurga containing lower diversity. Due to the emphasis on the presence or absence of fungal isolates and diversity, the statistical analysis is further constrained by the fact that it is not quantitative. Because of this, some fungal groupings are more likely to be found than others, independently of their quantity. Despite the fore mentioned drawbacks, our methodology is adequate for identifying variations in factor affecting fertility and fungal diversity across soils and environmental conditions. The present study is a proper evaluation of factors (abiotic and organic) influencing the role of fungal diversity affecting the soil fertility variations across two drastically diverse regions of Karnataka. Thus giving an understanding of the climatic and ecological conditions playing an important scientific role in modulating the microbial (fungal) diversity and also maintaining the bioavailability and recycling of nutrients. The detailed characterisation of the fungal species diversity, their biochemical properties and physiological role in colonizing the soil environments are in due course and would be the extended outcomes of this presently reported work. This would yield clear, detailed insights into the prominence of wildly adopted fungal species and their diversity in regulating bioconversion, bioavailability and recycling of nutrients affecting soil fertility.

